# The effects of parasitism on sex allocation of a hermaphroditic acorn barnacle

**DOI:** 10.1101/2024.04.20.590428

**Authors:** Masami M. Tamechika, Hiroyuki Yamada, Shigeho Ijiri, Yoichi Yusa

**Author notes:** **Corresponding author contact details:** Masami M. Tamechika.

## Abstract

Sex allocation theory predicts the adaptive allocation of resources to male versus female reproduction in simultaneous hermaphrodites in response to individual characteristics or environmental factors. Because parasites uptake resources from their hosts, their presence could affect the sex allocation of the hosts. We investigated the effects of infestation status and infestation intensity by the rhizocephalan barnacle *Boschmaella japonica* on reproduction, including sex allocation, of the host intertidal barnacle *Chthamalus challengeri*. Feeding activity was also examined as a factor related to resource intake. Both male and female reproductive investment decreased with increasing parasite infestation, and the sex allocation of large infested hosts was more male-biased than that of large uninfested hosts. Moreover, in contrast to the model prediction that male investment does not change under resource limitation, male investment decreased in infested hosts whose resources were taken by parasites. This reduction in male investment could be explained by changes in mating group size, since infested hosts have shorter penises and consequently are able to access fewer mating partners.

## Introduction

Sex allocation, defined as the allocation of resources to male versus female reproduction, is an important life-history trait that affects reproductive success in sexual organisms (Charnov 1982). Sex allocation is expressed as sex ratio in dioecious organisms, the timing and direction of sex change in sequential hermaphrodites, and resource allocation to male and female functions in simultaneous hermaphrodites (Charnov 1982; Schärer 2009; West 2009). The theory of sex allocation seeks to predict the optimal sex allocation for individuals acted on by natural selection, and it does so by considering the shapes of fitness gain curves of male and female functions for various individual traits or environmental conditions.

Simultaneous hermaphroditism is a common sexual system in plants and some animals (Jarne and Auld 2006). Simultaneous hermaphrodites are useful in studies of sex allocation, which is simply expressed as the optimal resource allocation to male versus female functions (Schärer 2009). Unlike dioecious organisms, in simultaneous hermaphrodites each individual expresses both sexual traits simultaneously (Michiels 1998), and hermaphrodites often exhibit flexible allocation (reviewed by Charnov 1982; Schärer 2009). Consequently, changes in sex allocation immediately affect fitness through reproductive success in one generation (Borgia and Blick 1981; Michiels 1998). Because of these characteristics, the effects of different individual traits and environmental conditions on sex allocation can be monitored simultaneously, and studies using simultaneous hermaphrodites to test the predictions of sex allocation theory have been conducted extensively (Schärer 2009).

Basic sex allocation theory for simultaneous hermaphrodites assumes that female fitness is not limited by sperm availability but by the amount of own reproductive resources, since the greater cost of producing a gamete for females (eggs) than males (sperm) limits female reproductive success (Bateman 1948; Charnov 1979; Anthes et al. 2010). Therefore, the female fitness gain curve does not show diminishing returns and increases almost linearly with resources invested. On the other hand, the male fitness gain curve in a small mating group shows diminishing returns with increasing resource investment since the number of eggs in the group is limited and sperm competition occurs (Fig. 1a; Charnov 1979, 1982). However, details may differ among organisms with different life histories and reproductive systems. For example, brood space is limited in brooding organisms, which leads to a saturation of female fitness (Fig. 1d; Heath 1977; Charnov 1982; p. 226; Strathmann et al. 1984). Sex allocation in this case is predicted to be more male-biased than in organisms without brooding limitations. For male fitness, many factors have been shown to alter the degree of saturation of the gain curve, such as the frequency of selfing (Johnston et al. 1998), the level of sperm competition (Petersen 1991), and, above all, mating group size (MGS) (e.g., Raimondi and Martin 1991; Trouvé 1999; Tan et al. 2004).

**Fig. 1.**
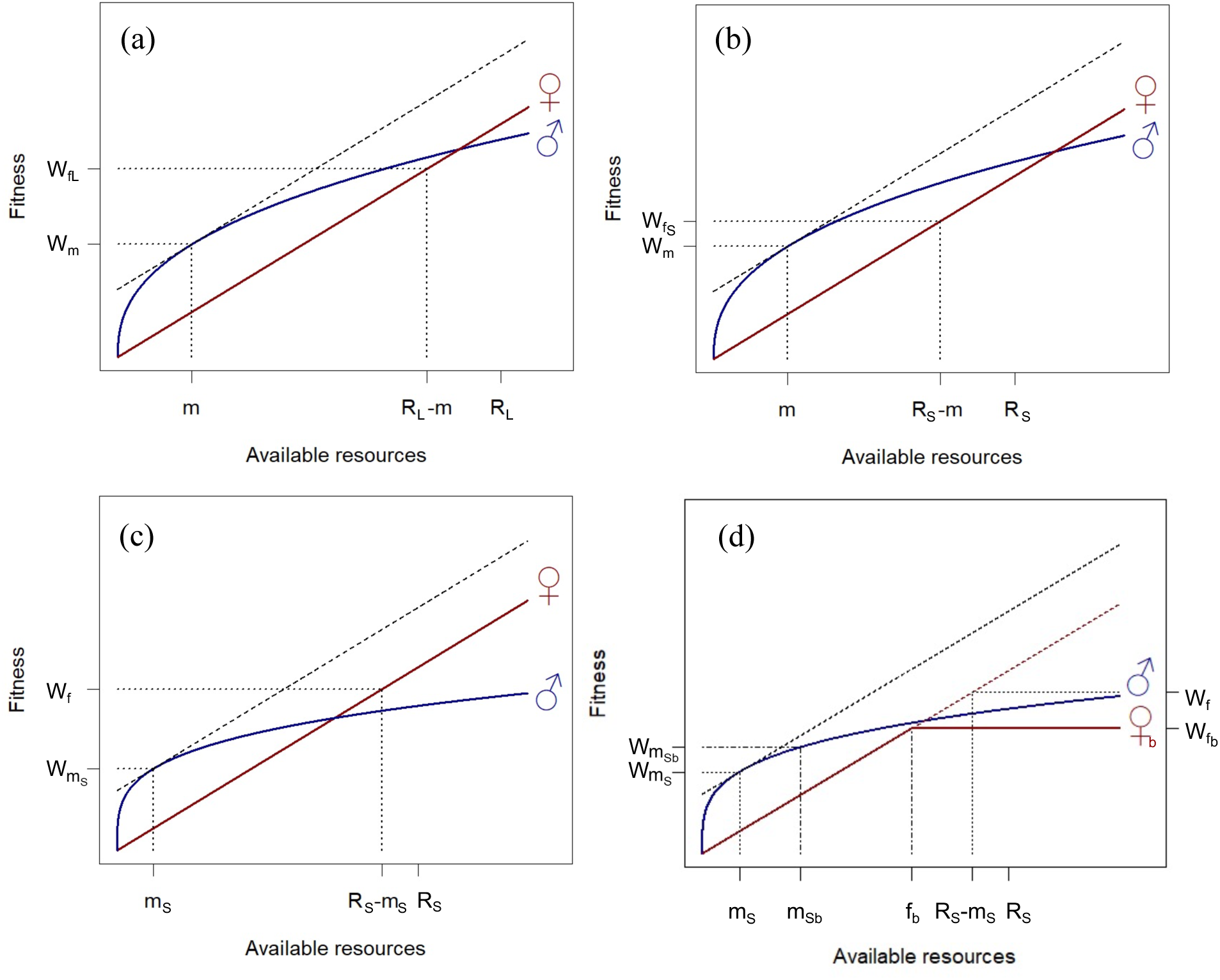
Hypothesized fitness gain curves for simultaneous hermaphrodites under (a) basic, (b) resource limitation, (c) small mating group size, and (d) brood space limitation in sex allocation theory. The female gain curve (red) is assumed to be linear, and the male gain curve (blue) is given by *W_m_ = r^n^* with diminishing returns due to sperm competition, where *W_m_* is male fitness, *r* is resource input, and *n* provides the various shape of the fitness gain curve in relation to mating group size. (a) The maximum fitness of individuals with *R_L_* resources is realized by allocating *m* to male function and the rest (*R_L_ − m*) to female function, where the rates of increase (slope) for both male and female gain curves are equal. (b) Under resource limitation (*R_S_* < *R_L_*), the optimal resource input *m* remains the same, but the female input is reduced to *R_S_ − m*. (c) Under small mating group size (small *n*), *m* is reduced to *m_S_* since the male gain curve rises steeply. (d) Under brood space limitation, the female input is reduced to *R_S_ – f_b_*. Accordingly, the male input increases as *R_S_ − (m_S_ + f_b_*). Modified from Frank (1987a, b), Charnov (1982, p. 224–226), Vizoso and Schärer (2007), Schärer (2009), and Hart (2010).

Although various individual or intraspecific traits have been explored as factors affecting sex allocation, the effects of interspecific relationships, such as predation or parasitism, on the sex allocation of hermaphrodites are largely unexplored (Schärer 2009). Among such interspecific relationships, parasites are especially interesting because they affect host life-history traits by depriving the host of energy and consequently changing its resource allocation patterns for growth and reproduction (Sorensen and Minchella 2001; Lafferty and Kuris 2009).

Parasites can be categorized into two groups in terms of their pathways of energetic influence on the host: consumers and castrators (Baudoin 1975; Hall et al. 2007). Consumers merely consume the host’s resources, whereas castrators both consume the host’s resources and directly affect the allocation of resources by the host, such as by affecting growth rate (Takahashi and Matsuura 1994; Sorensen and Minchella 2001; Miura et al. 2006) or investments into reproduction and sexual characteristics (Lafferty 1993a; Corral et al. 2021). Therefore, castrators may directly change the sex allocation (e.g., offspring sex ratio) of the host to maximize their own fitness, as has been reported for the bacterium *Wolbachia* in arthropod hosts (Werren et al. 2008). On the other hand, consumers may indirectly influence the resource allocation of the host, such as by affecting size at maturity (Lafferty 1993b; Jokela and Lively 1995; Fredensborg and Poulin 2006) or selfing rate (Morran et al. 2011). In such cases, hermaphrodites might adaptively shift their allocation pattern to minimize the effect of infestation on fitness (Schärer 2009).

Barnacles are suitable for studies of sex allocation as they have diverse life histories and sex allocation, with several species having pure males or females (Darwin 1854; Yusa et al. 2012, 2013; Lin et al. 2015). Sex allocation theory was originally developed for barnacles (Charnov 1980, 1982, 1987), and empirical studies have shown the adaptive plasticity of sex allocation in hermaphroditic barnacles (Raimondi and Martin 1991; Yusa 2018). Barnacles are sessile and mate using a long penis (Anderson 1993; except for the unique mode of “sperm casting”: Barazandeh et al. 2013, 2014; Barazandeh and Palmer 2015), and these characteristics facilitate the determination of MGS by researchers (Hoch 2008; Neufeld and Palmer 2008; Tamechika et al. 2020).

Some barnacles are infested by parasites such as isopods and rhizocephalans (Crisp 1960; Høeg et al. 1990; Høeg 1995; Fong 2016; Fong et al. 2019a). For example, the parasite *Boschmaella japonica* (Rhizocephala: Chthamalophilidae) (Deichmann and Høeg 1990; Høeg et al. 1990, 2019) is known to impede reproduction, but not completely castrate, its host intertidal barnacle *Chthamalus challengeri* (Thoracica: Chthamalidae) (Høeg et al. 1990; Yabuta et al. 2020). This parasite is likely a consumer as the parasite’s nutrient-absorbing organs, the internae, do not appear to damage host organs (Bresciani and Høeg 2001). Multiple sac-like reproductive organs, the externae, emerge from the mantle as well as the prosoma (body) of the host (Høeg et al. 1990). It is possible that the host barnacle adaptively changes its sex allocation in response to the resource limitation caused by the parasite in this system.

Here, we investigate the effects of *B. japonica* infestation status (presence or absence) and infestation intensity (number and total area of parasite externae) on the reproduction and sex allocation of its host barnacle. Additionally, the feeding activity of the host was assessed as a factor relevant to total resource availability, as infested individuals could conceivably overcome the effects of resource limitation by improving their food intake (Polak 1996).

## Methods

### Sampling

*Chthamalus challengeri* specimens were collected at mid-tidal level on a rocky shore in Shirahama, Wakayama, Japan (33°41′38″N, 135°20′15″E) in May 2019. The prevalence of *Boschmaella japonica* is high in April and May, and *C. challengeri* broods eggs from February to June at this site (Yabuta et al. 2020). We collected five boulders (length range 46–68 cm, mean 52.6 cm; width range 26–54 cm, mean 35.4 cm) each inhabited by 43–170 (mean 81) barnacles.

### Cirral activity

We examined the cirral activity of unparasitized and parasitized barnacles. We placed the five collected boulders individually in aquaria (21 × 30 × 5 cm high) with running natural seawater at the Seto Marine Biological Laboratory immediately after sampling. All barnacles were acclimated to laboratory conditions for at least 24 h. Then, each boulder was gently moved to another aquarium of the same size without water flow for video recording. We conducted behavioral records for 5 min by using a digital camera (EverioR, JVC, Yokohama Japan). Later, all barnacles were fixed in 99.5% ethanol and were checked for infestation status by dissection. We randomly selected 60 uninfested and 60 infested individuals, analyzed the video, and counted the number of cirral beats in 5 min for each individual (see Appendix Table 1 for details of sample sizes).

**Table 1.**
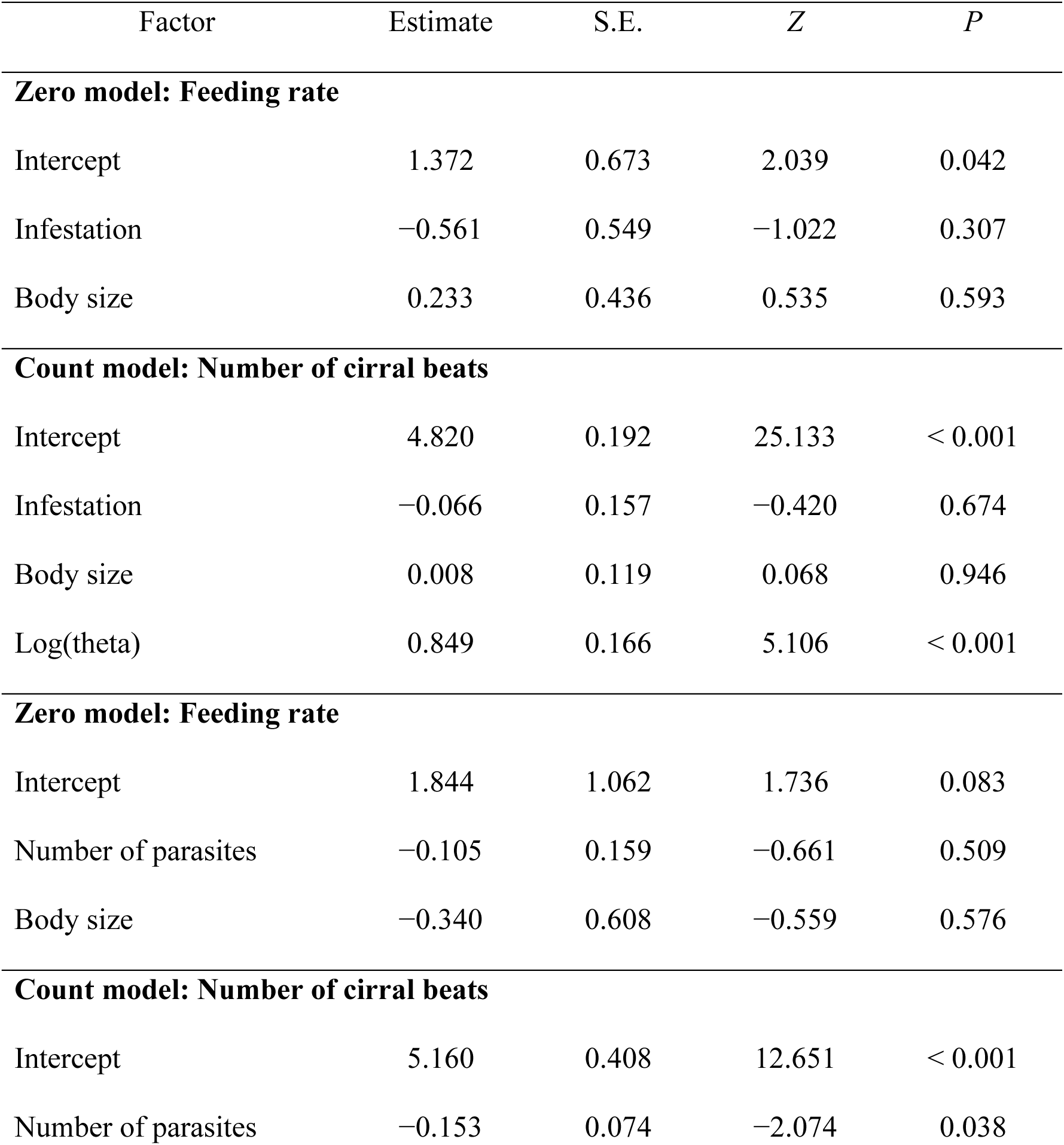
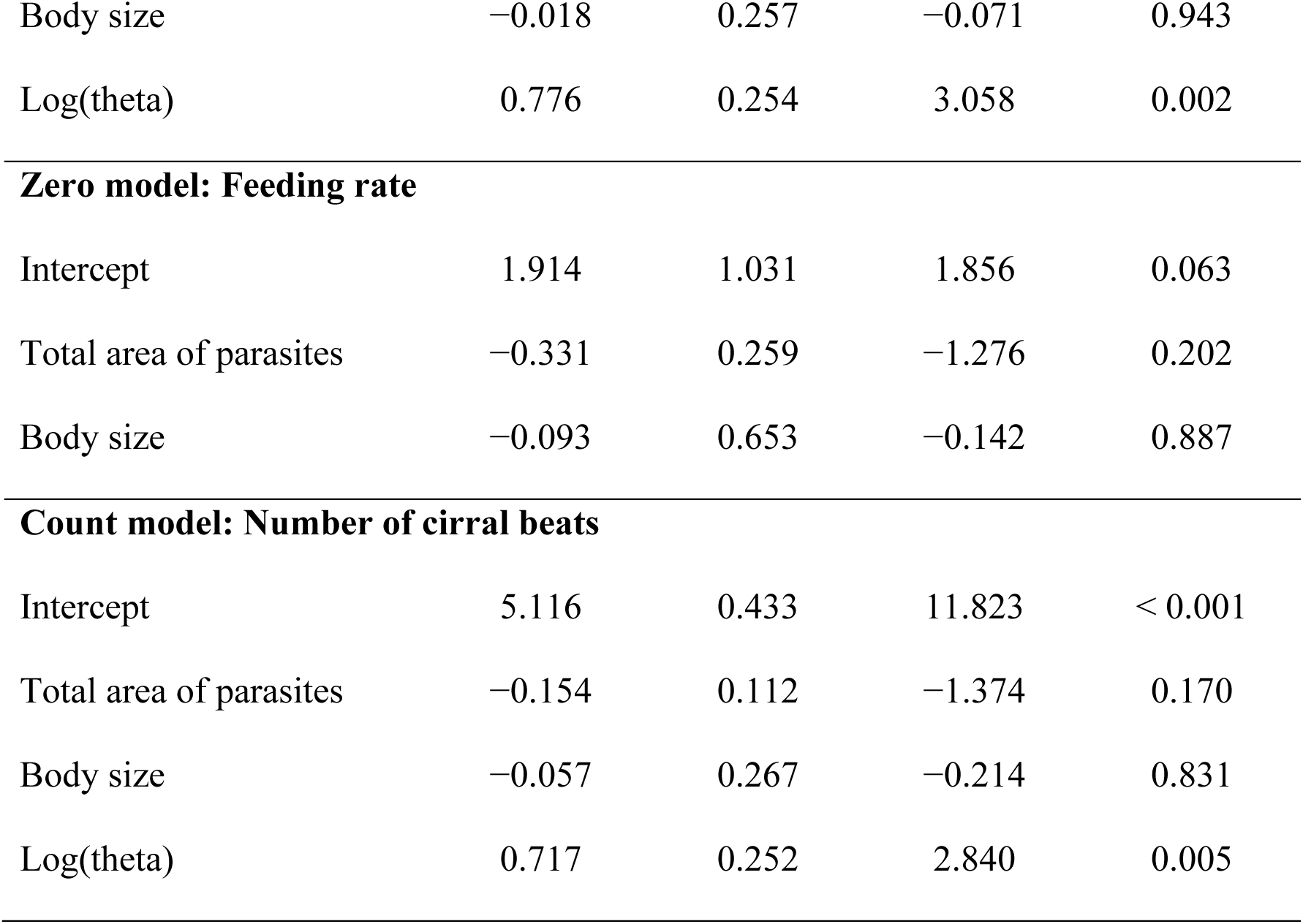
Hurdle model results on the effects of *Boschmaella japonica* infestation and host size on the behavior (feeding rate and number of cirral beats) of *Chthamalus challengeri*. Hurdle models with a binomial distribution and logit-link function were used for the zero component, and a negative binomial distribution and log-link function were used for the count data.

| Factor | Estimate | S.E. | Z | P |
| --- | --- | --- | --- | --- |
| **Zero model: Feeding rate** |  |  |  |  |
| Intercept | 1.372 | 0.673 | 2.039 | 0.042 |
| Infestation | −0.561 | 0.549 | −1.022 | 0.307 |
| Body size | 0.233 | 0.436 | 0.535 | 0.593 |
| **Count model: Number of cirral beats** |  |  |  |  |
| Intercept | 4.820 | 0.192 | 25.133 | < 0.001 |
| Infestation | −0.066 | 0.157 | −0.420 | 0.674 |
| Body size | 0.008 | 0.119 | 0.068 | 0.946 |
| Log(theta) | 0.849 | 0.166 | 5.106 | < 0.001 |
| **Zero model: Feeding rate** |  |  |  |  |
| Intercept | 1.844 | 1.062 | 1.736 | 0.083 |
| Number of parasites | −0.105 | 0.159 | −0.661 | 0.509 |
| Body size | −0.340 | 0.608 | −0.559 | 0.576 |
| **Count model: Number of cirral beats** |  |  |  |  |
| Intercept | 5.160 | 0.408 | 12.651 | < 0.001 |
| Number of parasites | −0.153 | 0.074 | −2.074 | 0.038 |
| Body size | −0.018 | 0.257 | −0.071 | 0.943 |
| Log(theta) | 0.776 | 0.254 | 3.058 | 0.002 |
| **Zero model: Feeding rate** |  |  |  |  |
| Intercept | 1.914 | 1.031 | 1.856 | 0.063 |
| Total area of parasites | −0.331 | 0.259 | −1.276 | 0.202 |
| Body size | −0.093 | 0.653 | −0.142 | 0.887 |
| **Count model: Number of cirral beats** |  |  |  |  |
| Intercept | 5.116 | 0.433 | 11.823 | < 0.001 |
| Total area of parasites | −0.154 | 0.112 | −1.374 | 0.170 |
| Body size | −0.057 | 0.267 | −0.214 | 0.831 |
| Log(theta) | 0.717 | 0.252 | 2.840 | 0.005 |

### Measurements

We checked for the presence or absence of parasites in the fixed barnacles by using a stereoscopic microscope. We counted the number of *B. japonica* externae and measured the area of the externae by using ImageJ (version 1.53k, National Institutes of Health). Then, we dissected out the operculum (consisting of two plates, the scutum and tergum), soma, penis, and eggs (if present) from the samples (see Appendix Table 1). The operculum was dried in an oven at 60 °C for 12 h and weighed to provide an index of body size (Kado et al. 2009; Tamechika et al. 2020). We counted the number of eggs for brooded individuals, and then randomly selected three egg capsules for each individual and measured their major and minor axes under a stereoscopic microscope. The volume of each egg capsule was calculated using these values by assuming an ellipsoid (Yabuta et al. 2020), and the three measurements were averaged for each brooded individual. Penis length, penis diameter, and the total volume of the testis and seminal vesicles were evaluated as male reproductive parameters. The penis was photographed with a digital camera (Stylus TG-4, Olympus Corporation, Tokyo, Japan), and the length and diameter were measured to the nearest 10 µm by using ImageJ. The volume of the testis and seminal vesicles was evaluated by sectioning the soma of randomly selected samples. The soma was fixed in 99.5% ethanol for 1 day for dehydration, embedded in paraffin, cut into 5-µm-thick sections with a microtome, and stained with hematoxylin and eosin. We photographed 10 sections at equal intervals from the beginning to the end of these organs, and then measured the area of the testis and seminal vesicles with ImageJ for each of the 10 sections. We used the next section if the selected section was damaged. The total volume of the testis and seminal vesicles was calculated as

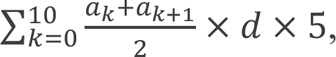

where *a_k_* and *a_k_*_+1_ are the average areas of the *k*th and *k* + 1th sections, *d* is the distance between adjacent sections, and *a* = 0 for both *a*_0_ and *a*_11_.

Next, we examined how MGS changes with penis length, which was estimated to be an average of 0.386 mm shorter for infested individuals than for uninfested ones (Table 2). We calculated the MGS for uninfested individuals and then recalculating the MGS with a decrease of 0.386 mm in penis length to approximate the MGS of infested individuals. The MGS was defined as 1 + the number of conspecifics reachable by the length of the elongated penis, *sensu* Charnov (1982). The number of reachable conspecifics was calculated from the distance between individuals (i.e., minimum distance between the two opercula) and the length of the elongated penis, which was assumed to be the normal penis length × 1.82, because, in the barnacle *Balanus grandula* on wave-protected shores, the penis elongates to 1.82 times the normal length at mating (Neufeld and Palmer 2008).

**Table 2.**
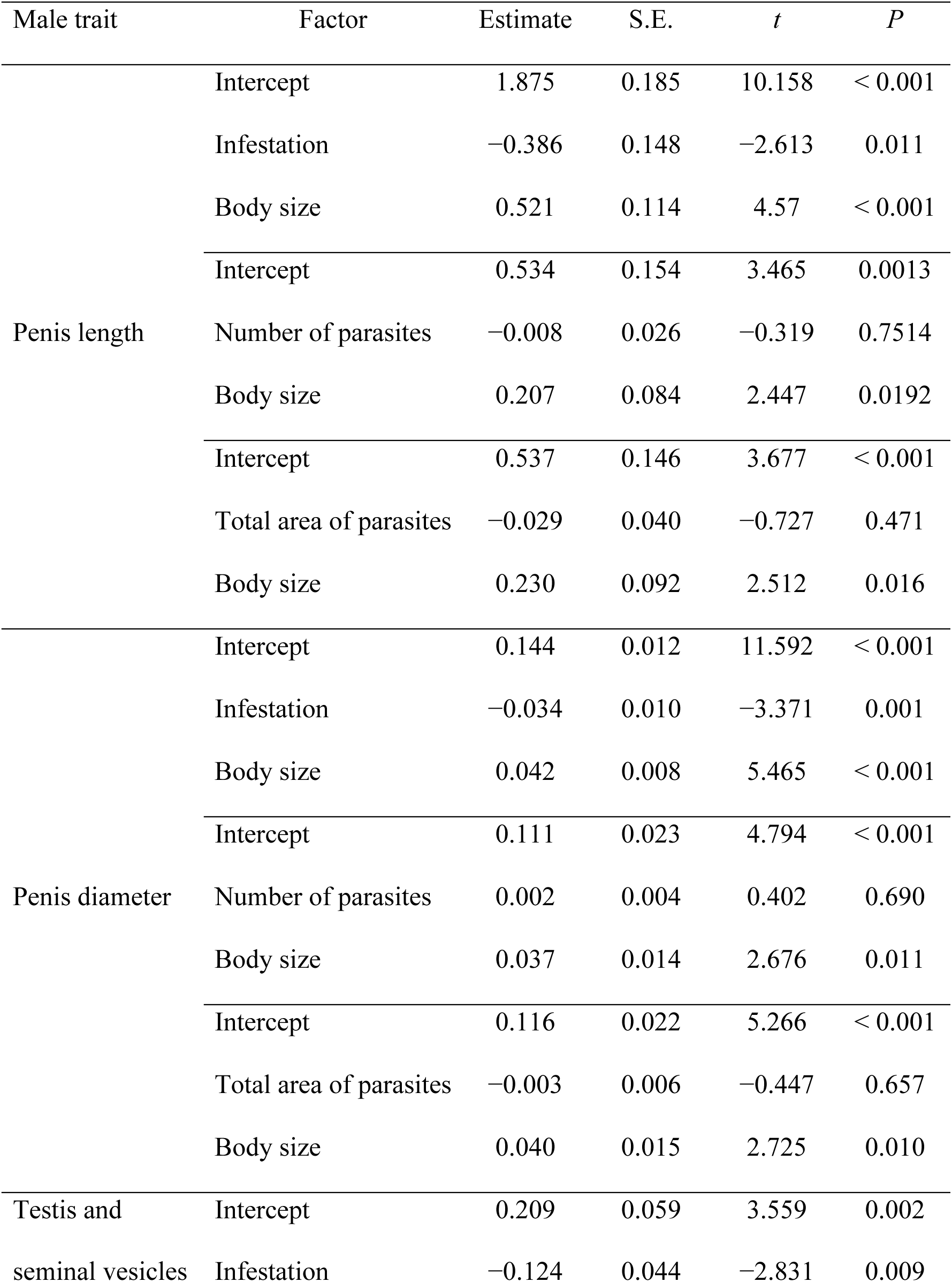
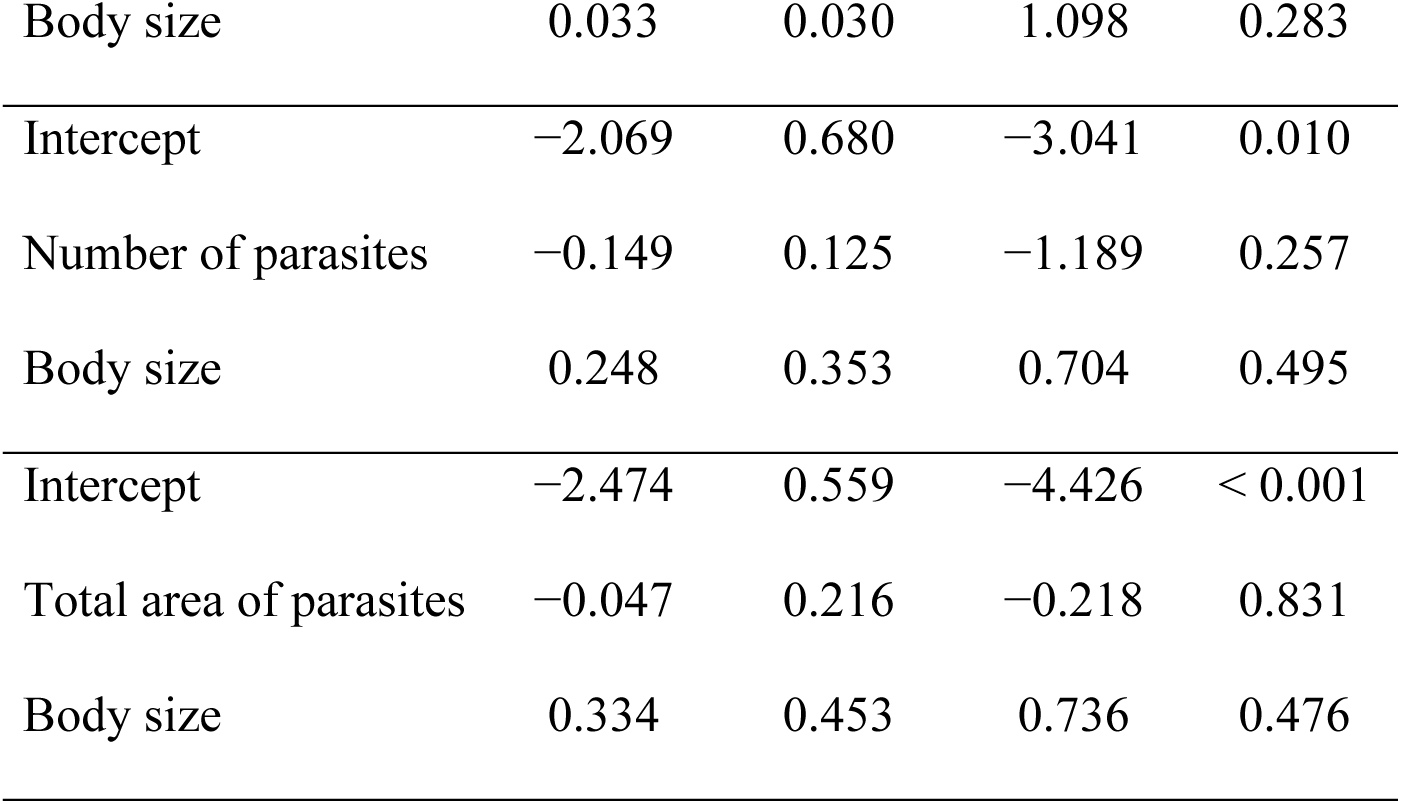
Generalized linear model results on effects of *Boschmaella japonica* infestation and host size on male reproductive investment of *Chthamalus challengeri*.

| Male trait | Factor | Estimate | S.E. | *t* | *P* |
| --- | --- | --- | --- | --- | --- |
| Penis length | Intercept | 1.875 | 0.185 | 10.158 | < 0.001 |
|  | Infestation | −0.386 | 0.148 | −2.613 | 0.011 |
|  | Body size | 0.521 | 0.114 | 4.57 | < 0.001 |
|  | Intercept | 0.534 | 0.154 | 3.465 | 0.0013 |
|  | Number of parasites | −0.008 | 0.026 | −0.319 | 0.7514 |
|  | Body size | 0.207 | 0.084 | 2.447 | 0.0192 |
|  | Intercept | 0.537 | 0.146 | 3.677 | < 0.001 |
|  | Total area of parasites | −0.029 | 0.040 | −0.727 | 0.471 |
|  | Body size | 0.230 | 0.092 | 2.512 | 0.016 |
| Penis diameter | Intercept | 0.144 | 0.012 | 11.592 | < 0.001 |
|  | Infestation | −0.034 | 0.010 | −3.371 | 0.001 |
|  | Body size | 0.042 | 0.008 | 5.465 | < 0.001 |
|  | Intercept | 0.111 | 0.023 | 4.794 | < 0.001 |
|  | Number of parasites | 0.002 | 0.004 | 0.402 | 0.690 |
|  | Body size | 0.037 | 0.014 | 2.676 | 0.011 |
|  | Intercept | 0.116 | 0.022 | 5.266 | < 0.001 |
|  | Total area of parasites | −0.003 | 0.006 | −0.447 | 0.657 |
|  | Body size | 0.040 | 0.015 | 2.725 | 0.010 |
| Testis and seminal vesicles | Intercept | 0.209 | 0.059 | 3.559 | 0.002 |
|  | Infestation | −0.124 | 0.044 | −2.831 | 0.009 |

|  |  |  |  |  |
| --- | --- | --- | --- | --- |
| Body size | 0.033 | 0.030 | 1.098 | 0.283 |
| Intercept | −2.069 | 0.680 | −3.041 | 0.010 |
| Number of parasites | −0.149 | 0.125 | −1.189 | 0.257 |
| Body size | 0.248 | 0.353 | 0.704 | 0.495 |
| Intercept | −2.474 | 0.559 | −4.426 | < 0.001 |
| Total area of parasites | −0.047 | 0.216 | −0.218 | 0.831 |
| Body size | 0.334 | 0.453 | 0.736 | 0.476 |

### Statistical analysis

All statistical analyses were performed with R software version 4.1.2 (R Core Team 2021). The effects of host body size and infestation on cirral activity, male reproductive investment (penis length, penis diameter, and the total volume of the testis and seminal vesicles), female investment (brooding rate, egg number, and egg capsule volume), and sex allocation were analyzed. The degree of infestation was evaluated in three ways: (1) the presence or absence of infestation; (2) the number of parasites (except for 0); and (3) the total area of externae on each host (infestation intensity). In general, generalized linear models (GLMs) were used for analyses. The error distributions and link functions were selected on the basis of AIC score, and when using a Gaussian distribution, we also checked normality with the Shapiro–Wilk test. Interaction terms were removed if the non-significance of the interactions was confirmed. In the following, we describe the details of the statistical models used for each analysis.

### Host size and behavior

To compare the body sizes of infested and uninfested barnacles, we used the exact Brunner–Munzel test implemented in the “lawstats” package (Joseph et al. 2023) in R, which does not assume normality. We also examined the effects of parasitism on host behavior (i.e., the occurrence and number of cirral beats) by using hurdle models implemented in the package “pscl” (Jackman 2020) to cope with the abundance of zero data that were not observation errors (Dicken and Booth 2013; Yamazaki and Koizumi 2017). The hurdle model had two components: logistic regression for zero or non-zero values (i.e., the occurrence of feeding), and negative binomial regression for count data (i.e., number of cirral beats).

### Male reproductive investment

The effects of parasitism on male investment were analyzed by using GLMs (Gaussian, identity-link). The effects of the number and total area of externae were analyzed by using GLMs for penis diameter (Gaussian, identity-link), penis length (Gaussian, log-link), and testis and seminal vesicle volume (gamma, log-link).

### Female reproductive investment

Brooding rate and the number of eggs were analyzed by using a hurdle model (brooding rate, binomial; number of eggs, negative binomial). The volume per egg capsule was analyzed by using a GLM (gamma, log-link).

### Sex allocation

Sex allocation was defined as male investment (i.e., the volume of testis and seminal vesicles) divided by total reproductive investment (i.e., the sum of the volumes of testis and seminal vesicles and eggs) (Vizoso and Schärer 2007). The total volume of eggs was calculated as the number of egg capsules multiplied by the egg capsule volume. We used the volumes of male and female gametes to calculate sex allocation because we do not know the actual energetic costs invested in each sexual function. This assumption has been used previously as a good proxy of relative sex allocation among individuals of the same species (reviewed in Schärer 2009; Janicke et al. 2013).

To test whether infestation and host body size affected sex allocation, we employed a GLM (gamma, log-link) with male investment as the response variable and total reproductive investment as the offset value (Nervo et al. 2014). Additionally, we checked the effect of body size on sex allocation for infested and uninfested individuals by using a GLM (gamma, log-link).

### Crowding effect

The crowding effect is the negative effect on parasites caused by competition among parasites within a host for finite resources (Read 1951; Fong et al. 2017). We checked whether host body size and crowding (i.e., the number of externae on the same host) affected the areas per externa and of all externae. The area per externa was analyzed by using generalized linear mixed-effects models (GLMMs) with Gamma distribution and log-link implemented with the packages “lme4” (Bates et al. 2015) and “lmerTest” (Kuznetsova et al. 2017). We incorporated the host barnacle’s ID as a random effect in this model. The total area of externae was analyzed by using a GLM (gamma, log-link).

## Results

### Host size and behavior

Host body size did not affect infestation status (i.e., presence or absence of parasites; Brunner–Munzel test, *P* = 0.066; uninfested 1.300 ± 0.103 mg; infested 1.497 ± 0.088 mg; mean ± S.E.; Appendix Fig. 2). The feeding rates of uninfested and infested individuals were 84% (*n* = 44) and 76% (*n* = 46), respectively, and the number of cirral beats per 5 min for uninfested and infested individuals ranged from 0 to 242 (105.36 ± 10.49) and 0 to 204 (92.80 ± 10.71), respectively. Neither infestation status nor host body size affected the number of cirral beats (Table 1).

**Fig. 2.**
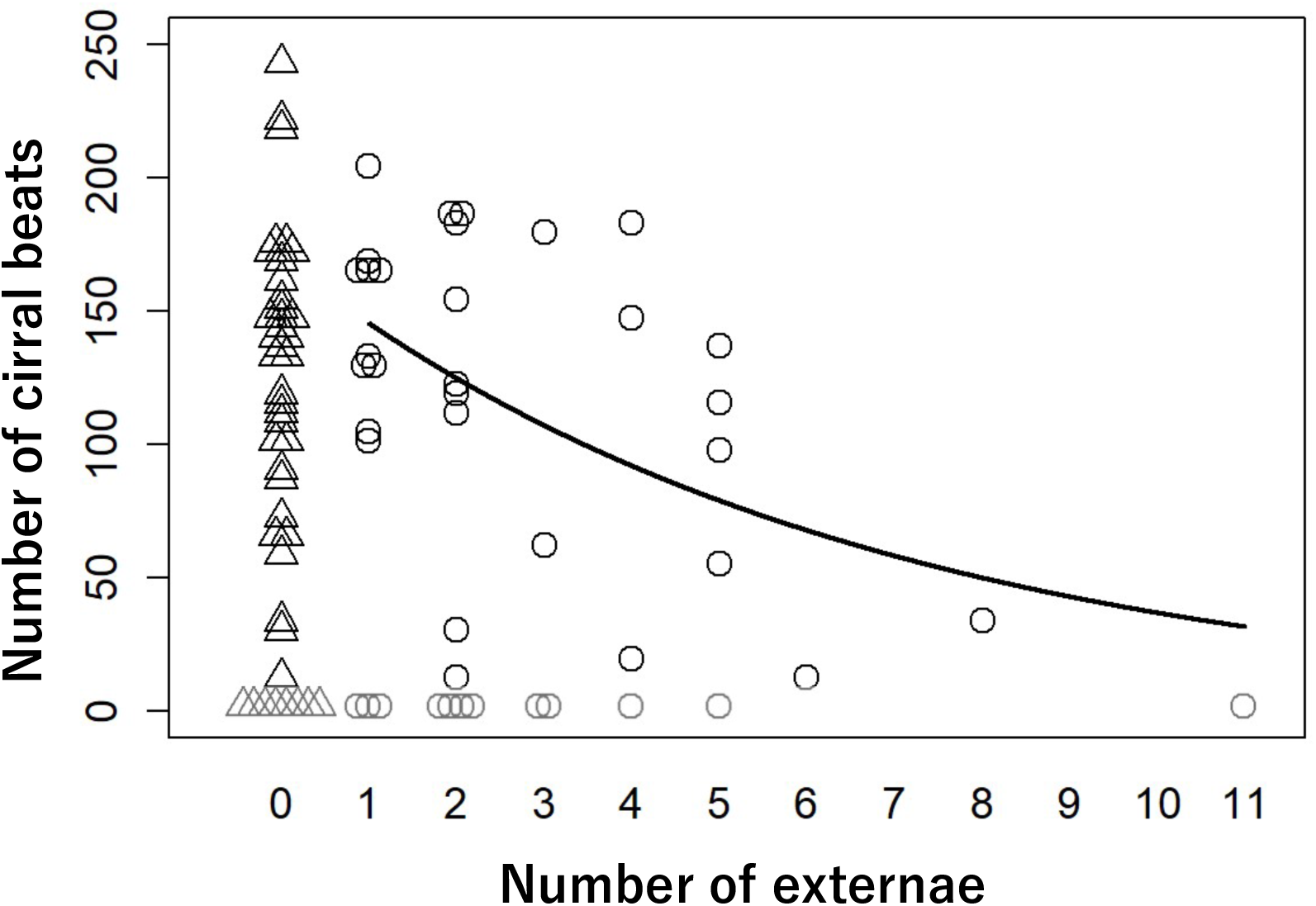
Effect of the number of externae of the parasite *Boschmaella japonica* on the number of cirral beats by the host barnacle *Chthamalus challengeri.* Triangles show uninfested barnacles, and circles show infested barnacles.

Among infested barnacles, the number of parasites ranged from 1 to 11, and the mean was 2.67 (Appendix Fig. 1). The total area of externae did not affect feeding rate or the number of cirral beats (Table 1). However, the number of cirral beats decreased as the number of parasites increased (Table 1; Fig. 2).

### Male reproductive investment

The averages and standard errors of penis length, penis diameter, and testis and seminal vesicle volume of uninfested individuals were 2.57 ± 0.11 mm, 0.20 ± 0.0078 mm, and 0.26 ± 0.039 mm^3^, respectively, and those of infested individuals were 2.27 ± 0.11 mm, 0.17 ± 0.0083 mm, and 0.13 ± 0.025 mm^3^. These three indices of male reproductive investment were significantly reduced by infestation (Table 2; Fig. 3). Penis length and diameter increased with body size, but testis and seminal vesicle volume did not change with size. We did not find any effects of the total area or number of parasites on all male reproductive investment (Table 2).

**Fig. 3.**
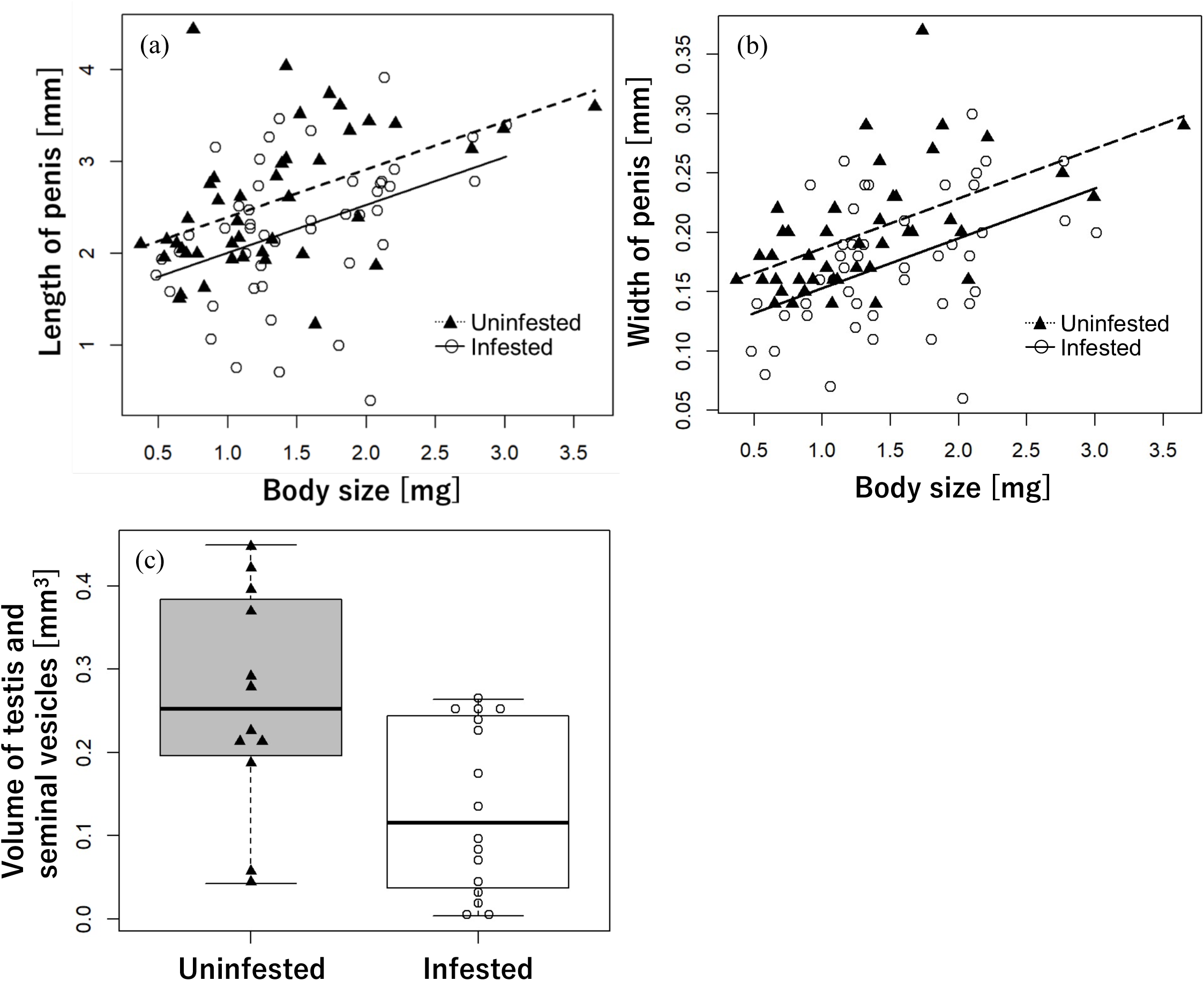
Effects of infestation by the parasite *Boschmaella japonica* on the host barnacle *Chthamalus challengeri* and host body size on male investment. (a) Penis length; (b) penis diameter; (c) volume of testis and seminal vesicles. Circles represent uninfested individuals; triangles represent infested individuals.

### Female reproductive investment

The brooding rate of uninfested individuals was 56% (*n* = 45) and that of infested ones was 33% (*n* = 46). Uninfested individuals produced 347.64 ± 58.99 eggs and infested ones produced 130.48 ± 37.05 eggs (mean ± S.E.). These indices of female reproductive investment were significantly reduced by infestation and increased by host size (Table 3; Fig. 4). However, the volume of per egg was not affected by either factor (Table 3; uninfested 6.66 × 10^−4^ ± 0.35 × 10^−4^ mm^3^, infested 7.39 × 10^−4^ ± 0.38 × 10^−4^ mm^3^; mean ± S.E.). Although the number of parasites had no effect on any indices of female reproductive investment, the total area of parasites affected the number of eggs (Table 3).

**Fig. 4.**
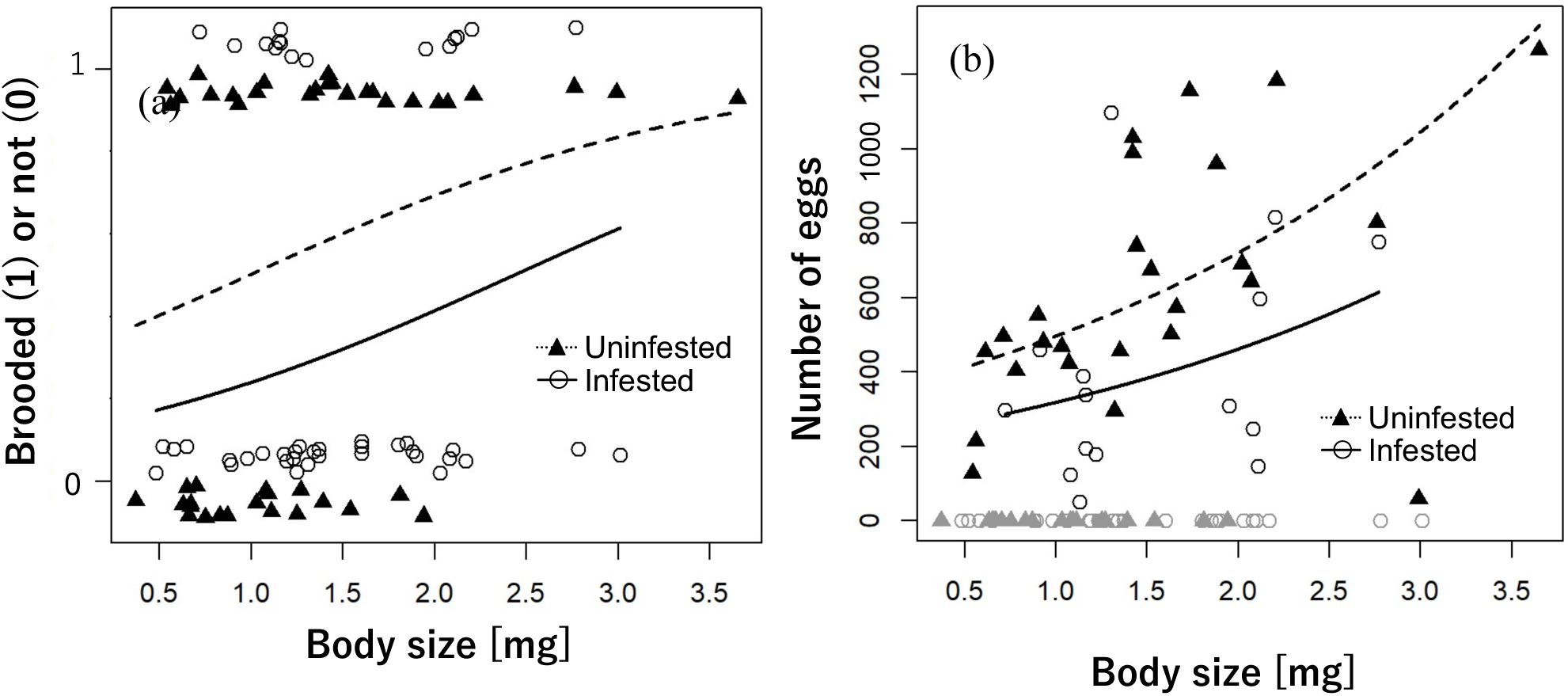
Effects of infestation by the parasite *Boschmaella japonica* on the host barnacle *Chthamalus challengeri* and of host body size on female investment. (a) Brooding rate; (b) number of eggs. Triangles and dotted lines represent infested individuals; circles and solid lines represent uninfested individuals.

**Table 3.**
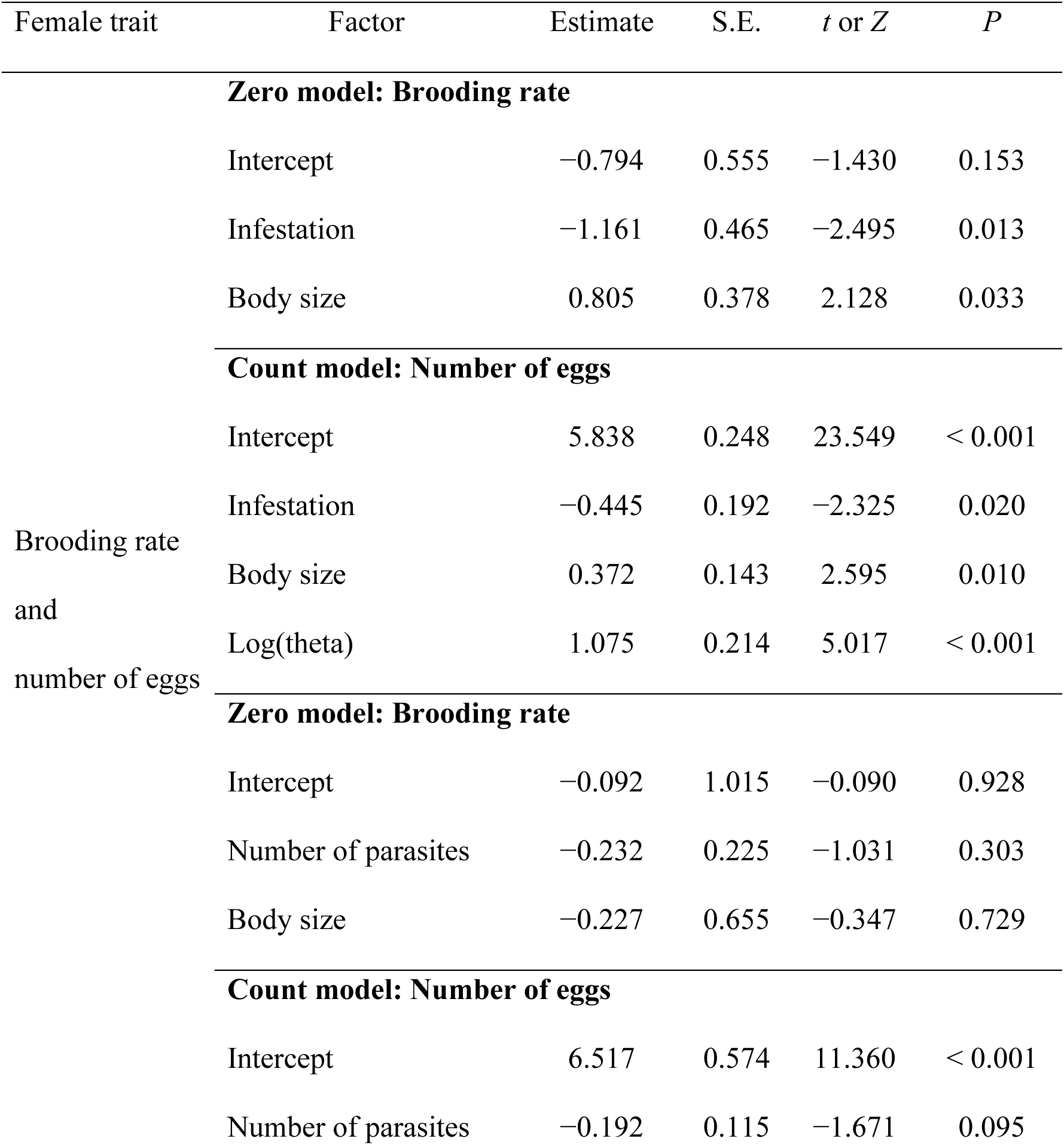
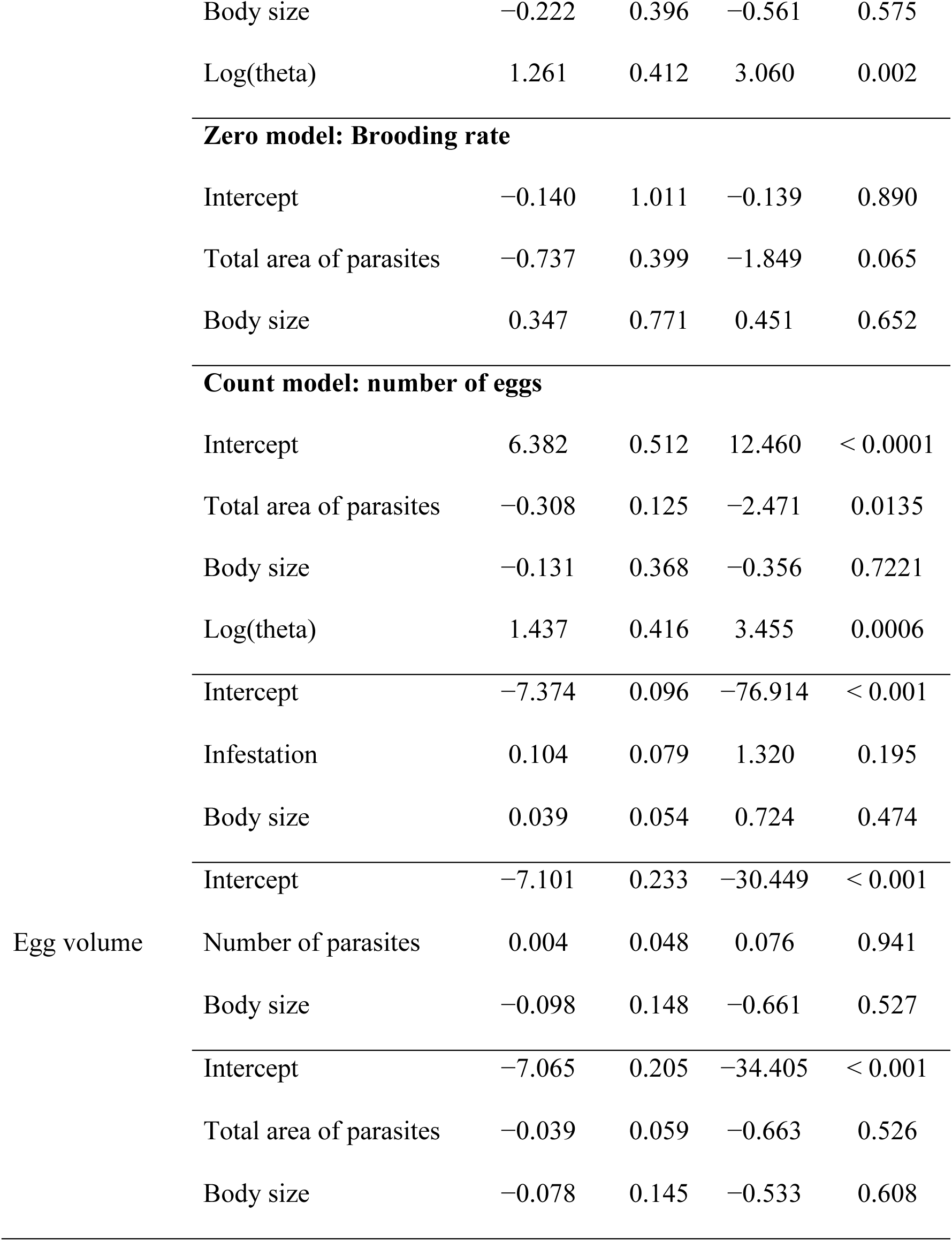
Hurdle model and generalized linear model results on effects of *Boschmaella japonica* infestation and host size on the female reproductive investment (brooding rate, number of eggs, and egg volume) of *Chthamalus challengeri.* Hurdle models with a binomial distribution and logit-link function were used for the zero component, and a negative binomial distribution and log-link function were used for the count data.

### Sex allocation

Host body size and the interaction between infestation and host body size significantly affected sex allocation (Table 4; Fig. 5). The number and total area of parasites did not affect sex allocation (Table 4). Sex allocation became more female-biased with increasing body size in uninfested individuals. However, sex allocation was not affected by body size in infested ones (Table 4).

**Fig. 5.**
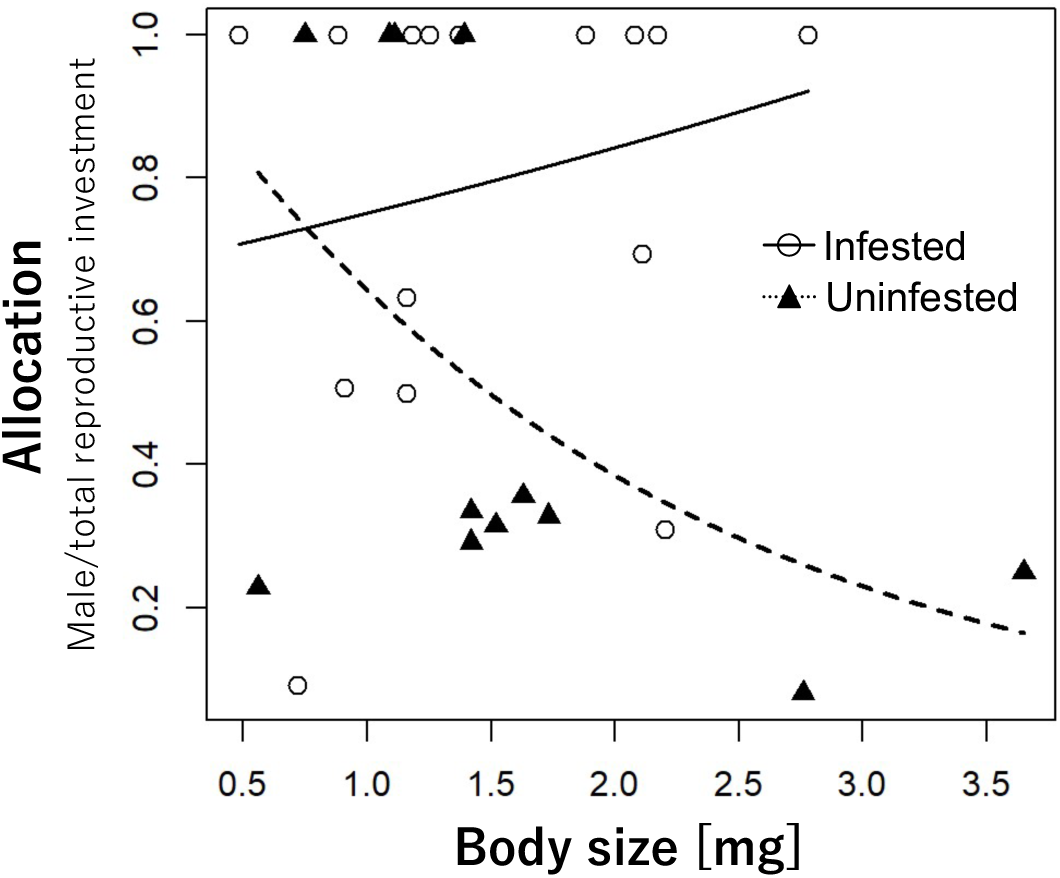
Effects of infestation by the parasite *Boschmaella japonica* on the host barnacle *Chthamalus challengeri* and of host body size on sex allocation (male reproductive investment divided by total reproductive investment). Triangles and dotted lines represent infested individuals; circles and solid lines represent uninfested individuals.

**Table 4.** Generalized linear model results on the effects of *Boschmaella japonica* infestation and host size on sex allocation by *Chthamalus challengeri*, and on the effects of host size on sex allocation by uninfested and infested individuals. Sex allocation was calculated as male reproductive investment divided by total reproductive investment. Generalized linear models were parameterized with a gamma distribution and log-link function.

| Target group | Factor | Estimate | S.E. | *t* | *P* |
| --- | --- | --- | --- | --- | --- |
| All individuals | Intercept | 0.075 | 0.310 | 0.244 | < 0.001 |
|  | Infestation | −0.477 | 0.442 | −1.078 | 0.292 |
|  | Body size | −0.516 | 0.174 | −2.972 | 0.007 |
|  | Infestation × body size | 0.631 | 0.262 | 2.407 | 0.024 |
|  | Intercept | −0.637 | 0.311 | −2.046 | 0.063 |
|  | Number of parasites | 0.044 | 0.057 | 0.768 | 0.458 |
|  | Body size | 0.224 | 0.161 | 1.388 | 0.190 |
|  | Intercept | −0.490 | 0.250 | −0.490 | 0.250 |
|  | Total area of parasites | 0.031 | 0.096 | 0.031 | 0.096 |
|  | Body size | 0.162 | 0.202 | 0.162 | 0.202 |
| Uninfested individuals | Intercept | 0.075 | 0.378 | 0.200 | 0.846 |
|  | Body size | −0.516 | 0.212 | −2.436 | 0.035 |
| Infested individuals | Intercept | −0.401 | 0.255 | −1.575 | 0.137 |
|  | Body size | 0.115 | 0.159 | 0.726 | 0.480 |

### Reduction of MGS

The MGS of uninfested and infested conditions ranged from 5 to 37 (11.46 ± 2.37, mean ± S.E.) and from 3 to 32 (8.85 ± 2.18), respectively. All individuals had a lower MGS with infestation, and the degree of reduction ranged from 1 to 5 (2.62 ± 0.37).

### Area of externae

The area per externa and of all externae ranged from 0.072 to 3.348 mm^2^ and from 0.212 to 7.319 mm^2^, respectively. The area per externa declined with increasing number of externae on the same host (Table 5). The area of all externae increased with host body size and the number of externae on the same host (Table 5).

**Table 5.** Generalized linear mixed-effects model results on the effects of crowding (number of *Boschmaella japonica* parasites per *Chthamalus challengeri* host) and host size on area per externa, and generalized linear model results on the effects of crowding and host size on the area of all externae. The generalized linear mixed-effects model was parameterized with a gamma distribution and log-link function, and the generalized linear model with a Gaussian distribution and log-link function.

| Parasite trait | Factor | Estimate | S.E. | d.f. | *t* | *P* |
| --- | --- | --- | --- | --- | --- | --- |
| Area per externa | Intercept | −0.449 | 0.330 | 107 | −1.363 | 0.173 |
|  | Number of parasites | −0.095 | 0.044 | 107 | −2.176 | 0.030 |
|  | Host body size | 0.299 | 0.186 | 107 | 1.604 | 0.109 |
|  | **Random effect** | **S.D.** |  |  |  |  |
|  | Host barnacle ID | 0.383 |  |  |  |  |
| Area of all externae | Intercept | −0.346 | 0.252 | 39 | −1.375 | 0.1769 |
|  | Number of parasites | 0.160 | 0.022 | 39 | 7.430 | < 0.0001 |
|  | Host body size | 0.368 | 0.112 | 39 | 3.295 | 0.0021 |

## Discussion

Our findings indicate that the sex allocation of the hermaphroditic barnacle *C. challengeri* was affected by infestation by the rhizocephalan parasite *B. japonica*. Both female and male reproductive investments by the host barnacle were reduced by infestation, and in large individuals sex allocation became more biased toward males. Additionally, the number of cirral beats, which is related to food intake, decreased as the number of parasites increased. Below, we discuss the effects of infestation on each trait and the possibility that the changes in sex allocation are a result of resource limitation, rather than any specific adaptations against the parasite.

Parasite infestation negatively affected female reproductive investment, including brooding rate and the number of eggs, a finding that is consistent with those of Yabuta et al. (2020). Resource uptake by the parasite appears to be the reason for the negative effects, although the cost of defense against the parasite may also be relevant (Hall et al. 2007). A similar decline in female investment has been reported in other hermaphrodites (e.g., Sorensen and Minchella 2001; Adamo 2002; Calado et al. 2005, 2006; Fong et al. 2019a, b). Additionally, the decrease in the number of eggs may have been caused by a limitation of brooding space, as both the parasites and host eggs occupy the mantle cavity (Deichmann and Høeg 1990; Høeg et al. 1990), and the number of eggs decreased with increasing total area of parasites.

Few studies have examined the effect of parasite infestation on male investment in simultaneous hermaphrodites, and the few studies that have done so reported inconsistent trends. For example, land snails infested by parasitic mites produced larger sperm than uninfested ones (Haeussler et al. 2014). However, infestation status by a parasitic isopod did not affect male reproductive success in a protandric simultaneous hermaphrodite, the caridean shrimp (Calado et al. 2005). In our study, infestation by the parasite affected male investment in *C. challengeri*. Male investment can be distinguished into two categories of cost (Heath, 1977; Schärer 2009): fixed costs refer to the production and maintenance of reproductive organs, such as the genitalia, that are not consumed in each mating event, and variable costs refer to the resources allocated to produce sperm, which is consumed in each mating event. To our knowledge, no studies have simultaneously explored the effects of parasites on fixed and variable costs in a hermaphroditic animal. In this study, we found that parasite infestation reduced both fixed costs (i.e., genitalia: penis length and diameter) and variable costs (i.e., sperm in the testis and seminal vesicles).

The MGS as males (i.e., the number of female partners within reach of the penis; Yamaguchi et al. 2013; Tamechika et al. 2020) is expected to be reduced by infestation, as the penises of infested barnacles are shorter than those of uninfested ones. In sessile animals such as barnacles that mate using a penis, MGS is limited by the area within reach of the penis (Neufeld and Palmer 2008; Hoch 2016). Therefore, the shorter penises of parasitized barnacles will directly reduce MGS as male. In addition, the movements of the penis are controlled by longitudinal muscles and hydraulic pressure in the haemocoel (Klepal et al. 1972; Anderson et al. 1988; Klepal 1990; Neufeld and Palmer 2008), and penis diameter correlates with cuticle and muscle mass (Neufeld and Rankine 2012). Thus, the thinner penis of infested individuals may have lower mobility than that of uninfested individuals and possibly result in even smaller MGS as male. This would reduce the potential mating success of males and might be a reason for the reduced male reproductive investment of infested individuals. In future studies, this reduction in MGS by infestation should be further examined through paternity analysis.

The sex allocation of unparasitized individuals became more female-biased with increasing body size. This trend likely reflects local sperm competition and the budget effect (*sensu* Klinkhamer et al. 1997; Cadet et al. 2004). Specifically, the total number of eggs in each mating group is limited, which causes intense sperm competition (Charnov 1980, 1982; Schärer 2009). Under these conditions, male fitness follows the law of diminishing returns with resource input, whereas female fitness increases almost linearly with input. Large individuals can use more resources than small ones (i.e., the budget effect), and hence in hermaphrodites it might be adaptive to increase the investment to female functions more than to male functions (Vizoso and Schärer 2007; Schärer 2009). Such size-dependent sex allocation has been reported in both model and empirical studies on many hermaphroditic animals (e.g., Petersen and Fischer 1996; Schärer et al. 2001; Baeza 2007; Vizoso and Schärer 2007).

However, the tendency toward size-dependent allocation disappeared and the allocation became more male-biased in large infested individuals than in large uninfested ones. This relatively more male-biased sex allocation in infested individuals can be explained by the reduction of reproductive resources due to parasitism (Fig. 1a, 1b). Infested individuals have fewer total reproductive resources to allocate than do uninfested individuals. This means that infested individuals have limited resources available for reproduction irrespective of body size, and in this respect resemble small uninfested individuals. Under such conditions, the optimal sex allocation will be more male-biased (Fig. 1b, point *m*) because the rates (slopes) for both male and female gain curves are identical, as in small individuals. In fact, both models and empirical studies show that resource limitation leads to more male-biased sex allocation (Charnov 1982; Frank 1987; Vizoso and Schärer 2007). However, the present result slightly but clearly differs from the predictions of optimal sex allocation models under resource limitation. Whereas the models predict that investment in male functions remains constant with resource input (constant male hypothesis: Yamaguchi 1985; Frank 1987), the present results show that male investment also decreased. We suggest that this is related to the lower MGS of infested individuals due to the reduction of penis size (the leftward shift of point *m* to *m_S_* in Fig. 1b, 1c). In addition, the limitation of brooding space by parasite externae might limit fitness gains through female functions and thus favor a shift in allocation toward male functions (Fig. 1d, point *m_Sb_*).

We consider that this male bias in sex allocation is not a result of specific adaptation to the parasite, since the host *C. challengeri* is distributed widely from Hokkaido, Japan, to the East China Sea (Utinomi 1949, 1970; Kim and Kim 1980; Southward and Newman 2003; Cheang et al. 2012) and has a long planktonic larval phase (approximately 12–16 days; Chen et al. 2024), whereas the parasite *B. japonica* is only known to occur at three sites (Aburatsubo and Jogashima, Kanagawa, and Shirahama, Wakayama, Japan; Høeg et al. 1990; Ogawa and Matsuzaki 1991; Yabuta et al. 2020). It is therefore unlikely that adaptive plasticity of sex allocation to parasites has been maintained in this parasite–host system, and the shift in sex allocation caused by infestation is more likely a general response to resource limitation in this barnacle.

We found a negative effect of the number of parasites on the number of cirral beats, which was likely caused by limitation of the movement of the prosoma by parasites attached to various places on the host. Because barnacles feed by cirral movement (Crisp and Southward 1961), this could have a negative impact on food intake and thus reproductive budget. However, the number and total area of parasites did not affect other host traits such as brooding rate, male reproductive investment, or sex allocation. This might reflect a crowding effect among the parasites, as the number of parasites on the same host negatively affected the area per parasite. A similar crowding effect on parasite growth rate and fecundity has been reported for the rhizocephalan barnacle *Clistosaccus paguri* (Høeg 1982) as well as for other parasites (Read 1951; Roberts 2000; Heins et al. 2002; Valero et al. 2006; Fong et al. 2017). Therefore, an increase in the number of parasites might reduce the effect of each parasite on its own.

In conclusion, our study reveals the effects of parasite infestation on sex allocation by a hermaphroditic host. Infestation reduced both female and male reproductive investment in a simultaneously hermaphroditic barnacle. Moreover, whereas uninfested barnacles had a female-biased allocation with increasing body size, infested barnacles did not follow this trend, and large infested barnacles had a more male-biased allocation than large uninfested barnacles. These findings shed new light on the evolution of optimal sex allocation patterns under parasitism and, more broadly, on the complicated effects of species interactions in natural ecosystems.

## Author contributions

MT and YY designed the experiment; MT collected and analyzed the data with help from YY, SI, and HY; MT and YY led the writing of the manuscript, and all authors contributed to later versions and gave final approval for publication.

## Acknowledgments

We are grateful to Shigeyuki Yamato and the other members of the Seto Marine Biological Laboratory, Kyoto University; Ryoko Odake, Musashi Kitagawa, and the other members of the Fish Reproductive Physiology and Plankton Laboratory of Hokkaido University; Noriko Yasuoka of the Research Institute of Environment, Agriculture and Fisheries, Osaka Prefecture; Ryo Nakayama of the Fisheries Research Institute, Aomori Prefectural Industrial Technology Research Center; and Haruka Ootuki of Sakura Finetek Japan Co., Ltd., for their help with laboratory procedures. We also thank to Satoshi Wada and the other members of the Laboratory of Animal Ecology, Hokkaido University, for their helpful discussion on this manuscript. This study was supported by a JSPS KAKENHI grant (no. 19H0328), a JST SPRING grant (no. JPMJSP2119), and by the Research Institute of Marine Invertebrates Foundation.

## Data Accessibility Statement

All data that support this article will be posted to the Dryad repository as soon as this paper is published.

## Conflict of interest

The authors declare that they have no conflicts of interest.

## Notes

### Competing Interest Statement

The authors have declared no competing interest.

