## Supplemental 3 figures and 1 tables for "The effects of parasitism on sex allocation of a hermaphroditic acorn barnacle"

**Appendix Figures**

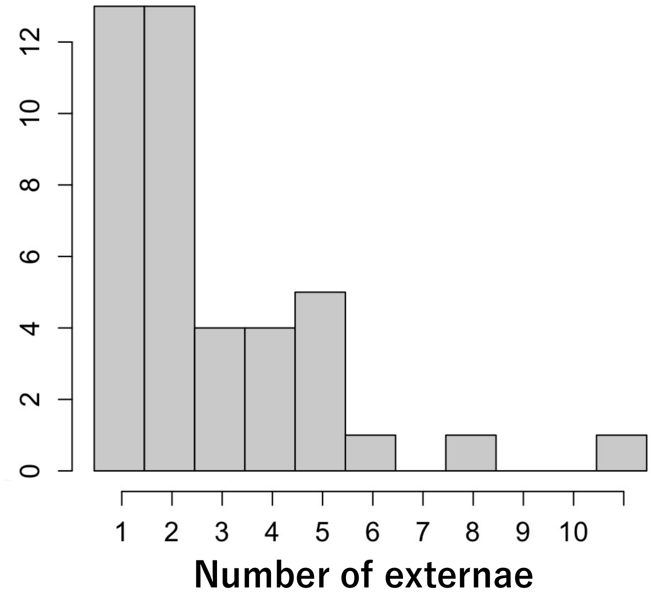


**Appendix Fig. 1.** Histogram of the number of externae among infested individuals. Infestations of one host by multiple parasites was observed, and the maximum number of parasites per host was 11.

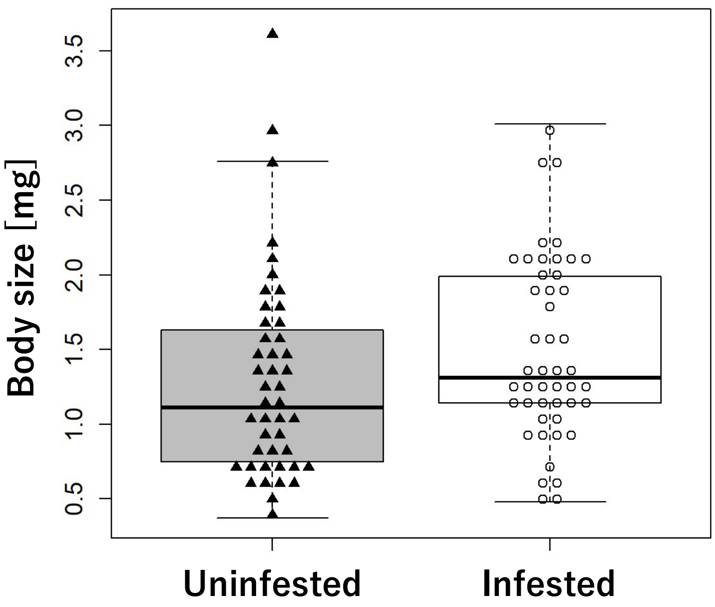


**Appendix Fig. 2.** Relationship between parasite infestation and body size. Triangles show infested individuals; circles show uninfested individuals.

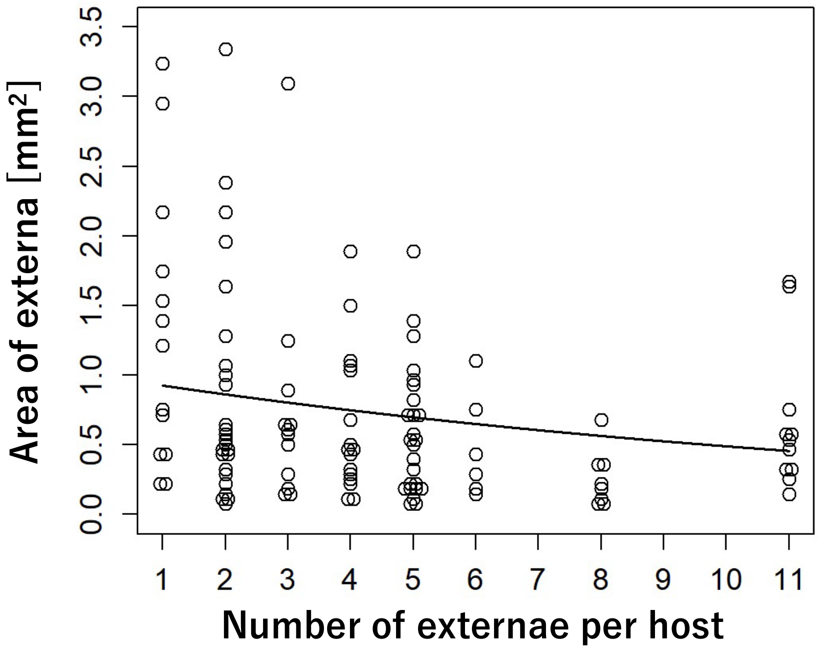


**Appendix Fig. 3.** Relationship between the area per externa and the number of externae per host.

### Appendix Tables

**Appendix Table 1** Full list of sample sizes, missing data, and reasons for data exclusion.

| | | | Uninfested | Infested | Missing | | Used for analysis | | Reasons for exclusion |
|---|---|---|---|---|---|---|---|---|---|
| | | | | | Uninfested | Infested | Uninfested | Infested | |
| First random selection | | | 60 | 60 | - | - | - | - | |
| Body size (operculum) | | | 60 | 60 | 15 | 13 | 45 | 47 | Partly broken |
| Parasites | | Number of externae<br>Volume of externae | - | 47 hosts | - | 5 hosts | - | 116 parasites in 42 hosts | Partly broken |
| Cirral activity | | Infestation status | 45 | 47 | 1 | 1 | 44 | 46 | Unavailable for the observation |
| | | Number of externae<br>Volume of externae | - | 47 | - | 5 | - | 42 | |
| Male function | Penis length and diameter | Infestation status | 45 | 47 | 2 | 1 | 43 | 46 | Partly broken |
| | | Number of externae<br>Volume of externae | - | 47 | - | 6 | - | 41 | |

| | | | | | | | | | |
|---|---|---|---|---|---|---|---|---|---|
| | | Infestation status | 40 | 40 | 28 | 24 | 12 | 16 | Mistakes in making serial sections |
| | Volume of testis and seminal vesicles | Number of externae<br>Volume of externae | - | 40 | - | 25 | - | 15 | Mistakes in making serial sections and the externae are partly broken |
| Female functions | Brooding rate and number of eggs | Infestation status | 45 | 47 | 0 | 1 | 45 | 46 | Partly broken |
| | | Number of externae<br>Volume of externae | - | 47 | - | 6 | - | 41 | |
| | Volume of egg capsules | Infestation status | 25 | 16 | 0 | 1 | 25 | 15 | |
| | | Number of externae<br>Volume of externae | - | 16 | - | 5 | - | 11 | |
| Mating group size | | | 45 | - | 32 | - | 13 | - | Unavailable for observation |
| Sex allocation | | Infestation status | 12 | 16 | 0 | 0 | 12 | 16 | - |
| | | Number of externae<br>Volume of externae | - | 16 | - | 1 | - | 15 | Partly broken |
